# Hypoglossal motor output is altered by C4 epidural electrical stimulation via ascending spinal and peripheral feedback pathways

**DOI:** 10.64898/2026.04.01.715924

**Authors:** Alyssa Mickle, Jesús Peñaloza-Aponte, Caitlin Brennan, Erica A Dale

## Abstract

After cervical spinal cord injury (cSCI), swallowing dysfunction is common and increases mortality via aspiration pneumonia. While these deficits have often been attributed to secondary damage from complications of injury management, there has recently been a greater appreciation for the modulatory role of spinal populations in swallow generation that are disrupted by injury. Here, we illustrate in a rodent model of cSCI that epidural spinal stimulation (ESS) of the phrenic motor nucleus at spinal segment C4 alters motor output at the hypoglossal motor nucleus through activation of excitatory ascending spinal pathways and inhibitory peripheral sensory feedback mechanisms. These findings highlight the importance of spinal-brainstem communication in shaping the motor program of swallow-related musculature and offer the potential for stimulation of the cervical spinal cord to be a therapeutic target for restoring swallowing function after injury.

**NEW & NOTEWORTHY:** In two varying severity models of spinal cord injury, we demonstrate the effects of spinal cord stimulation at C4 on the distal hypoglossal motor nucleus. We show that despite being anatomically distant, electrical stimulation of the phrenic motor nucleus increases hypoglossal motor output through ascending spinal pathways and dampens it through peripheral pathways. These findings highlight the importance of spinal-brainstem communication and illustrate the ability of spinal stimulation to restore this communication after injury.

## INTRODUCTION

After cervical spinal cord injury, swallowing dysfunction, also known as dysphagia, occurs in as many as 40% of patients and increases risk of aspiration pneumonia (Shem, Castillo et al. 2011). Dysphagia post injury is often attributed to damage of nearby peripheral nerves, anatomical alterations, consequences of spinal surgery, intubation, and weakness of respiratory muscles rather than interruption of neural communication between spinal and brainstem nuclei (McRae, Morgan et al. 2023). However, there is known coordination of breathing and swallow motor output in the form of Schluckatmung, or concurrent diaphragm activation during swallow thought to generate negative intra-thoracic pressure to position a bolus into the esophagus (Pitts and Iceman 2023). Indeed, there is increased recognition of the role of ascending spinal pathways in forming swallow patterns and their suppression by cervical spinal cord injury (Pitts, Iceman et al. 2022), though the precise nature of spinal inputs coordinating primarily swallow and primarily respiratory motor pools remains unclear.

Investigational clinical studies show that epidural spinal stimulation (ESS), or the subthreshold electrical stimulation of the dorsal surface of the spinal cord, restores a variety of motor functions after cervical spinal cord injury, particularly for locomotion (Lin, Shaaya et al. 2022). Beyond locomotion, we have previously shown in rats that ESS also restores patterned respiratory output after injuries that spare at least some residual connections to the brainstem respiratory central rhythm generators (Mickle, Penaloza-Aponte et al. 2024; Mickle, Penaloza-Aponte et al., *in review*). While ESS is typically applied below the lesion at the level of the targeted motor system’s spinal motor pool, there is evidence that electrical stimulation can also influence motor pools beyond the one being targeted. Within the respiratory circuit, ventral spinal pacing of intercostals at the thoracic spinal cord can concurrently activate the diaphragm via ascending spinal reflex pathways after a C2 complete transection (DiMarco and Kowalski 2013). This change in motor output from regions distal to stimulation highlights the potential for electrical stimulation to restore appropriate ascending spinal communication. Still, the ability for electrical stimulation to restore communication between motor pools below and above the injury remains unknown.

Here, we show that electrical stimulation of the spinal cord at C4 after a C2 hemisection injury increases motor output from the distal brainstem hypoglossal motor nucleus through diffuse neural circuitry and can transiently disrupt respiratory pattern generation. Completely severing ascending spinal pathways via C1 transection abolishes these excitatory effects and in turn leads to suppression of genioglossus electromyography (EMG) activity when stimulation occurs during the inspiratory phase. These data further illustrate the importance of spinal feedback in modulating accessory respiratory motor output and highlight the role of disrupted spinal-brainstem communication in swallowing difficulties after injury. Further applications of electrical stimulation may benefit by considering optimization of stimulation paradigms or electrode placement not just for facilitating top-down control of motor output, but also for restoration of key communications from the spinal cord to cortical and brainstem regions.

## METHODS

### Animals

All experimental procedures were approved by the Institutional Animal Care and Use Committee at the University of Florida and were conducted in accordance with NIH Guidelines Concerning the Care and Use of Laboratory Animals. This study is a secondary analysis of data derived from a subset of 6 rats previously described (Mickle, Penaloza-Aponte et al., *in review*). No additional experimental procedures were performed.

### Surgical preparation

Initial anesthetic induction was achieved via nosecone administration of 3% isoflurane in 40% O_2_ balance N_2_. After the implantation of femoral catheters, rats were converted to 0.17 mg/kg urethane anesthesia through a venous catheter. Reaction to toe pinch was regularly checked to ensure adequate depth of anesthesia. Two 36 AWG perfluoroalkoxy-insulated seven-stranded stainless-steel wire (AM Systems) EMG recording electrodes were implanted at the base of the tongue (targeting the genioglossus muscle), and bilaterally on the diaphragm. 40 AWG FEP-insulated stranded stainless-steel wires (Cooner wire) stimulating electrodes were implanted bilaterally on the dorsal surface of C4 by suturing to the dura. Electrodes were fabricated and implanted as described in detail previously (Holmes, Penaloza-Aponte et al. 2025). After electrode implantation, rats were cycle-triggered ventilated (AVS-1, CWE) using moving averaged genioglossus EMG to deliver breaths during respiratory effort in real time. After reaching steady-state on the mechanical ventilator, rats were right side C2-hemisected and given a 15-minute waiting period post-injury to allow for reaching steady-state levels of blood pressure and EMG amplitude before undergoing the first round of expiratory phase, inspiratory phase, and open-loop stimulation as detailed below. Rats were then completely transected at C1, given another 15-minute stabilization period, and stimulation repeated. Throughout the experimental period, respiratory frequency was allowed to fluctuate as dictated by the central control of the rat, and tidal volume of breaths was adjusted to maintain blood gases (see Mickle, Penaloza-Aponte et al., *in review*).

### Stimulating and recording parameters

Genioglossus EMG was sampled at 25 kHz, amplified (Model 1700 Differential AC Amplifier, AM Systems), band pass filtered (10-5000 Hz), notch filtered to remove 60 Hz powerline noise, digitized (Power1401-3A, Cambridge Electronic Design Limited) and recorded (Spike2 v10, Cambridge Electronic Design Limited). Electrical stimulation consisted of a biphasic, symmetric, charge-balanced, cathode leading 0.2 ms/phase square wave pulse delivered by an analog stimulus isolator (AM Systems Model 220, 1 mA/V). For inspiratory phase stimulation, a single stimulus pulse was delivered each time the raw genioglossus EMG signal crossed a threshold set at ∼75% the signal’s maximum amplitude at baseline. For expiratory phase stimulation, the pattern in which stimulation would have been delivered during a breath was recorded and then applied between genioglossal bursts. Open-loop stimulation was delivered at a constant 100 Hz throughout the respiratory cycle. Stimuli began at 50 µA in amplitude and increased by 50 µA steps to a maximum current amplitude of 250 µA. Each stimulus current amplitude was presented for ∼30 genioglossal bursts, after which there was a 1-minute rest period before presentation of the next current amplitude. After each round of stimulation from 50-250 µA was completed, there was a 5-minute washout period between each stimulation phase type (inspiratory, expiratory, open-loop).

### Data and statistical analysis

EMG signals were analyzed using custom MATLAB R2023b scripts available at github.com/dale-lab/ABRAP_Detangling. First, the raw genioglossal signal was enveloped using root-mean-square (RMS) with a 50 ms time constant and EMG burst onset/offset identified with an automated algorithm (Mickle, Rana et al. 2026). For each identified burst, EMG amplitude and area under the curve were calculated and normalized as fold change from the 30 seconds just prior to stimulation. Temporal coordination of EMG activity and stimulation was evaluated via a stimulus triggered average of the 25 ms directly post stimulus for each current pulse of a given amplitude and phase. GraphPad Prism 9 was used for statistical analysis. 2 and 3-way ANOVAs with the Geisser-Greenhouse correction of EMG response with current amplitude, EMG peak phase, stimulation phase, and/or injury type were conducted, with values of p < 0.05 considered statistically significant.

## RESULTS

After hemisection, the genioglossus responded directly to C4 electrical stimulation by increasing motor output, while this response was abolished after transection (**Figure 1**). That is, during inspiratory phase stimulation, the inspiratory phase genioglossus EMG RMS peak of hemisected but not transected rats was elevated (main effect injury, F(1, 5) = 11.78, p = 0.0186; interaction current and injury, F(1.523, 7.617) = 6.919, p = 0.0237; **Figure 2A**) as was expiratory phase genioglossus EMG during expiratory phase stimulation (main effect injury, F(1, 5) = 8.856, p = 0.0309; **Figure 2B**). In contrast to prior work in the diaphragm (Mickle, Penaloza-Aponte et al., *in review*)), expiratory phase electrical stimulus did not significantly increase inspiratory peak EMG response in either transected or hemisected rats (no main effect current, F(1.798, 8.992) = 3.304, p = 0.0872; no interaction current x injury, F(1.628, 8.140) = 2.063, p = 0.19; **Figure 2C**). While EMG peak was elevated in hemisected rats, after transection EMG output was suppressed specifically in response to inspiratory stimulation (main effect stimulation phase, F(0.3733, 1.867) = 21.57, p = 0.0447; **Figure 2D**). Also in contrast to the diaphragm (Mickle, Penaloza-Aponte et al., *in review*), genioglossus EMG did not remain elevated beyond the stimulation period after hemisection (**Figure 2E**) nor transection (**Figure 2F**).

**Figure 1:**
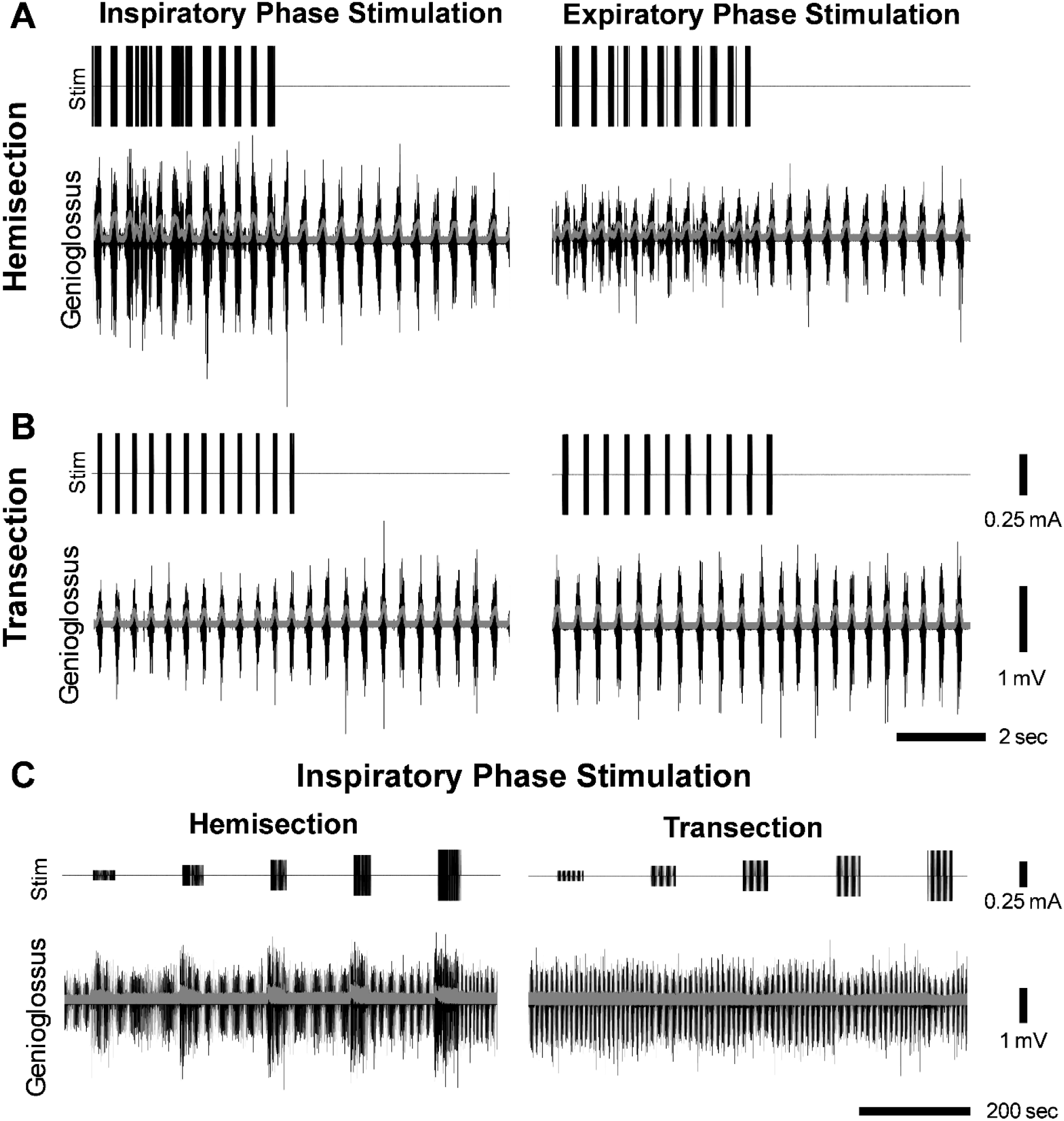
Representative traces of genioglossus EMG response to stimulation before and after C1 transection. Genioglossus EMG response to the highest amplitude 250 µA inspiratory and expiratory phase stimulation in A) C2 hemisected rats and B) transected rats. Traces are taken from the end of the stimulation period when EMG response was at ‘steady state’. C) Condensed trace of genioglossus EMG response to inspiratory phase stimulation across all stimulation levels showing increase in EMG after hemisection, and suppression of EMG at the highest currents after transection. For all traces - top, stimulation trace; bottom, genioglossus EMG (black: raw, gray: RMS envelope).

**Figure 2:**
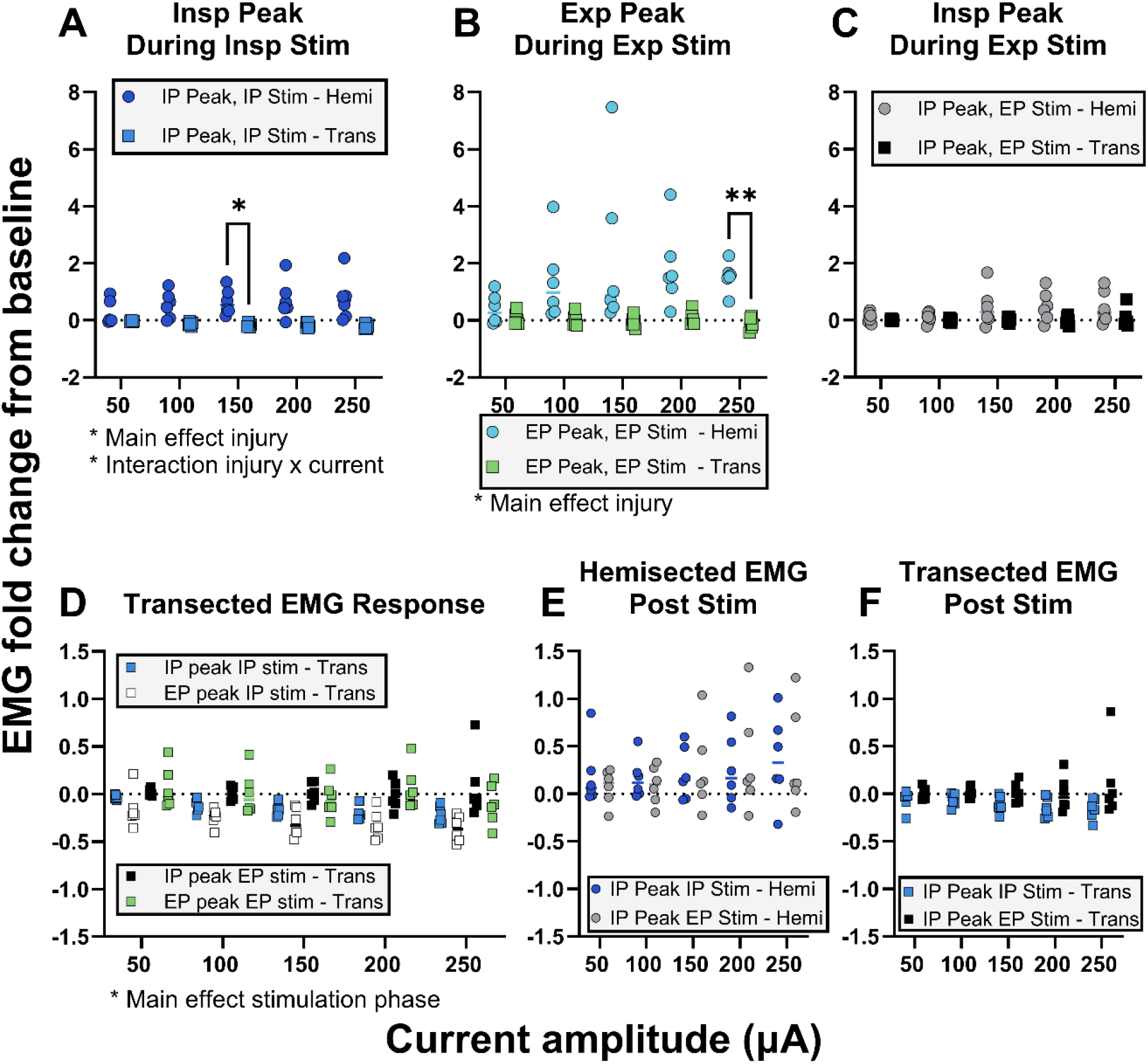
Hemisected rats increase genioglossus EMG in response to C4 stimulation during but not after stimulus, while inspiratory phase stimulation suppresses genioglossus EMG in transected rats. Change in genioglossus A) inspiratory phase EMG peak in response to inspiratory phase stimulation, B) expiratory phase EMG peak in response to expiratory phase stimulation, and C) inspiratory phase EMG peak in response to expiratory phase stimulation. D) Genioglossus EMG response to electrical stimulation in transected rats. Change in genioglossus inspiratory phase EMG peak of the 30 breaths immediately post stimulation in E) hemisected and F) intact rats. IP = Inspiratory phase; EP = Expiratory Phase; Hemi = C2 hemisection; Trans = C1 transection

Genioglossus EMG responses to current varied throughout the stimulation period. During the first few seconds of open-loop stimulation at the highest current amplitude, genioglossus EMG increased while contralesional hemidiaphragm activity was suppressed, causing short apneas in 3/6 rats, as seen in the representative trace (**Figure 3A**). In these rats, despite the suppression of contralesional activity, the ipsilesional hemidiaphragm continued to respond to stimulation. Even in those rats that did not experience disruptions in respiratory rhythm generation in response to open-loop stimulation, genioglossus EMG response was always highest in the first seconds of stimulation before tapering off. After transection, there was no increase in genioglossus EMG, which remained stable throughout the stimulation period (**Figure 3B**).

**Figure 3:**
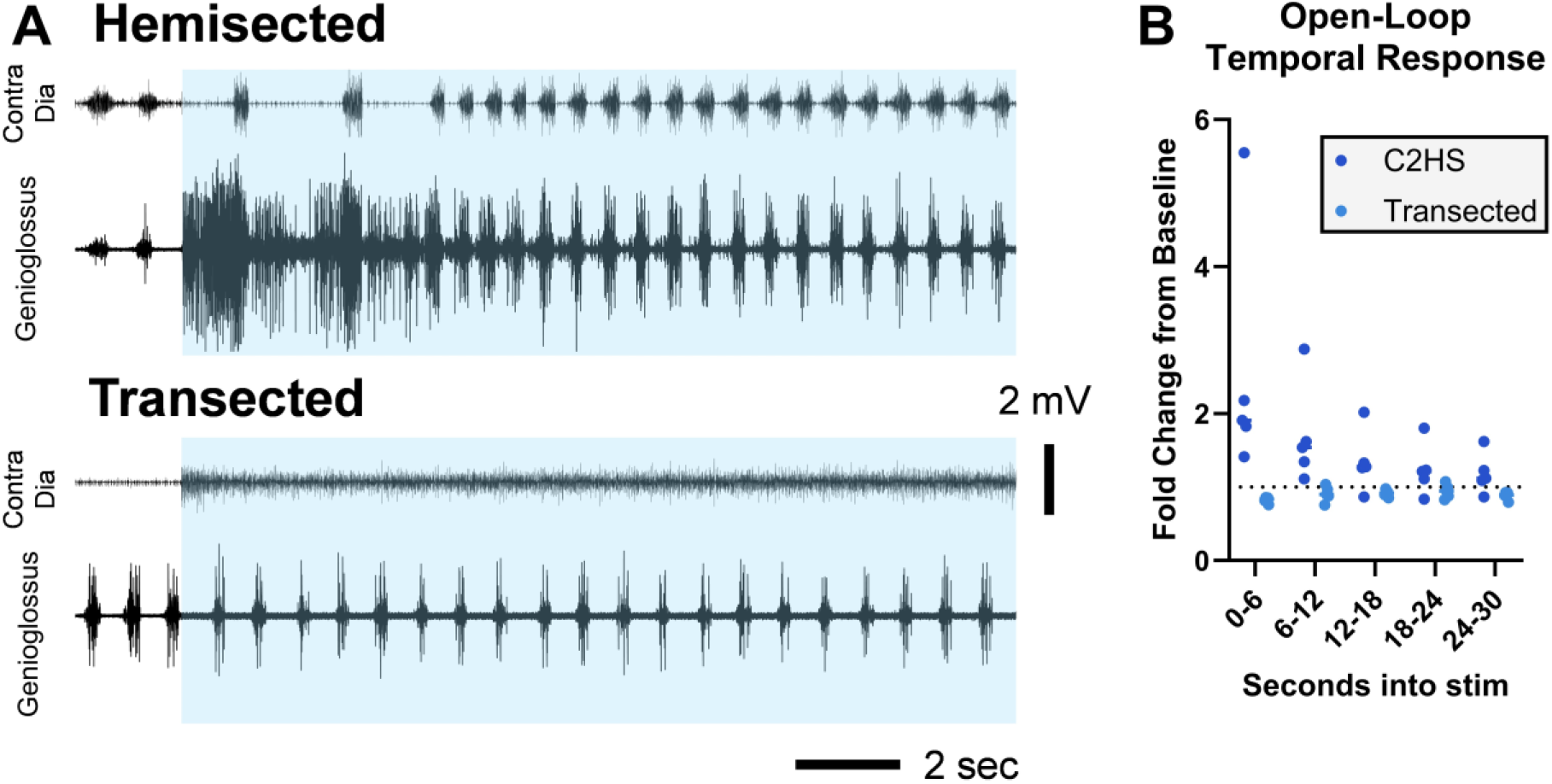
Initial response to stimulation can include respiratory disruptions which subside quickly. A) Representative traces of genioglossus EMG and respiratory disruptions caused by open-loop stimulation. Blue bars indicate time that 250 µA 100 Hz open-loop stimulation was on. B) Mean fold change of the peak genioglossus EMG RMS envelope during stimulation relative to the 30 seconds pre-stimulation.

As the latency of increased likelihood of EMG activation post-stimulus can give insights into the neural circuits at play, we next examined the temporal coordination of genioglossus EMG response to stimulus. Unlike with diaphragm EMG, increased firing likelihood at certain time points post stimulus was not immediately clear, particularly during inspiratory phase stimulation where EMG response is dominated by endogenous signal not related to stimulation (**Figure 4A**). However, upon closer examination, there was an increase in the likelihood of EMG signal particularly within 6-9 ms post stimulation during expiratory phase stimulus without the confound of endogenous firing. After transection, the 6-9 ms coordination was abolished, and stimulus related artifact primarily found in the early time bins post stimulation dominated the dampened response (**Figure 4B**). This temporal coordination of the genioglossus is much lower in magnitude than that of the ipsilesional diaphragm, indicating a more diffuse circuit is activated to attain increases in EMG, and occurs ∼2 ms delayed from the diaphragm response (**Figure 4C**).

**Figure 4:**
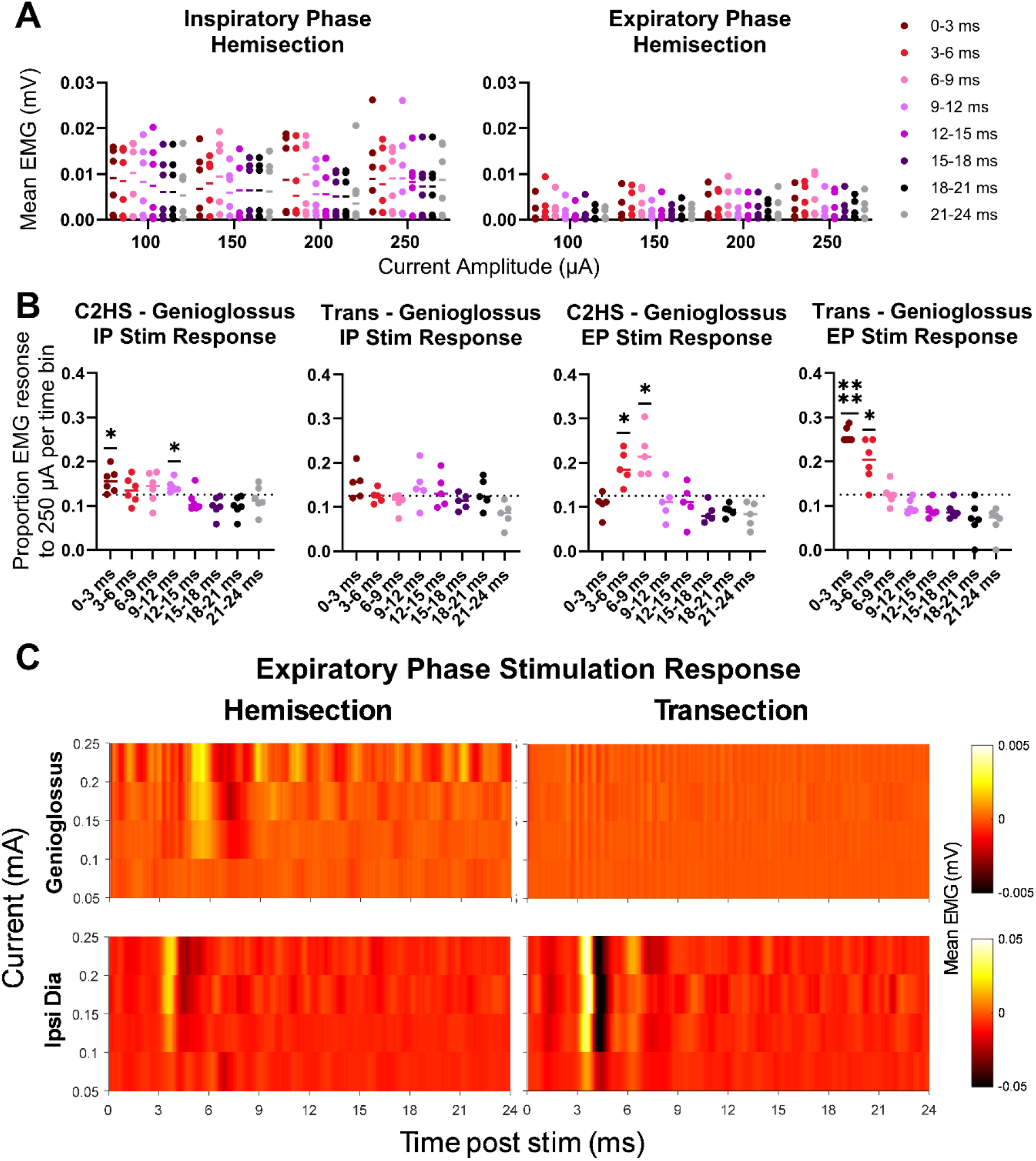
Genioglossus EMG shows weak temporal coordination to electrical stimulus. A) Mean stimulus triggered average of EMG response of the genioglossus to inspiratory and expiratory stimulation for the first 24 ms post stimulus. B) Proportion of the total mean stimulus triggered average EMG response is attributable to each time bin. Dashed line at 0.125 represents expected value if response was consistent across time bins. One sample t-tests used to determine what time bins EMG response proportion was significantly higher than 0.125. * p < 0.05, **** p < 0.0001. C) Sample representative top-down views of averaged genioglossus (top) and ipsilesional diaphragm (bottom) EMG response to expiratory phase stimulation traces after C2-hemisection (left) and C1-transection (right). Note the order of magnitude difference in scale of mean EMG values for genioglossus vs ipsilesional hemidiaphragm response to stimulation.

## DISCUSSION

Here we find that after a C2 hemisection injury, electrical stimulus at dorsal C4 increases hypoglossal motor output. This excitation of genioglossus EMG is mediated via ascending spinal pathways, as after transection, inspiratory phase stimulation suppresses genioglossus EMG output. Before transection, at the onset of high amplitude stimulus, respiratory pattern is disrupted with suppression of contralesional phasic diaphragm EMG, while the genioglossus continues responding with a tonic increase in EMG output. Together, these data suggest that electrical stimulation of the phrenic motor network not only restores diaphragm EMG output after injury but also facilitates bidirectional communication between the phrenic and hypoglossal motor nuclei.

### Potential neural substrates

As they share common anatomical substrates, swallowing and breathing must be highly coordinated behaviors. Breathing modulates swallow by establishing swallow timing, as most swallows occurring during expiration in healthy adults; in turn, swallow alters respiratory rhythm by inducing apnea as the bolus travels in the pharynx (Martin, Logemann et al. 1994). Beyond coordination of timing between the two behaviors, many components of the upper airway must change their activity between swallow and breathing. For example, during breathing, phasic activity of the genioglossus counters inspiratory pressure to prevent upper airway collapse (Tsuiki, Ono et al. 2000) while during swallow the genioglossus works to move material from the oral cavity to the pharynx (Kennedy, Kieser et al. 2010). This coordination of timing and switching of activity patterns of shared anatomical substrate is thought to be made possible via remodeling of activity within the brainstem networks controlling these behaviors (Davenport, Bolser et al. 2011), which can be accomplished by changing the activity patterns of neurons that are active during both behaviors (Saito, Ezure et al. 2003), or the way in which neural components of each behavior interact (Bolser, Poliacek et al. 2006). As such, there are many potential possibilities of neural substrates to allow for alteration of genioglossus EMG using stimulation targeted to the respiratory system.

The dorsal swallow group, a major component of central pattern generation for swallow, resides in the nucleus of the solitary tract (Jean 2008). For respiration, this region is also referred to as the dorsal respiratory group, a critical hub for processing sensory information related to pulmonary stretch reception (Ezure 2001). Importantly, sensory reporting of pulmonary stretch reception is delivered to the NTS peripherally via the vagal nerve. Increased pulmonary stretch reception may thereby be responsible for the decrease in genioglossus EMG after transection, which is by necessity a peripheral mechanism. Increased pulmonary stretch fits particularly with the stimulus phase-dependent nature of genioglossus EMG suppression, which occurred only during inspiratory stimulus. This inspiratory stimulus was paired with ventilation such that the diaphragm EMG induced by stimulus stacked with pulmonary stretch induced by the mechanical ventilation beyond that which occurred by ventilation alone at baseline. Additionally, stimulus during the inspiratory phase occurred concurrently with genioglossus EMG drive, indicating that this suppression is a transient phenomenon.

How precisely the ascending excitatory spinal signals recruited by stimulation at C4 before transection interact with brainstem centers to increase genioglossus EMG is less clear. The modest temporal coordination of genioglossus EMG to stimulation relative to the level of increase in total EMG activity indicates they may operate through a diffuse network to increase motor output. This may be reflective of the complex nature of interactions between the two pattern generators, with ascending spinal information traveling to and influencing output of multiple brainstem regions related to swallow and breathing. These regions then work together to increase genioglossus EMG output on differing timescales, and this diffuse nature of activation may further explain the changing the nature of diaphragm/genioglossus coordination at different current amplitudes.

At high current amplitudes after hemisection, the genioglossus had a greater initial response to electrical stimulation. This sometimes corresponded to suppression of phasic activity in the contralesional hemidiaphragm for the first seconds of stimulus. The shutting down of diaphragm EMG activity occurred only on the contralesional side and was not observed post-transection, implying active supraspinal suppression of breathing during this period. Indeed, breathing is inhibited at the level of brainstem control centers during swallow, even after laryngectomy (Hiss, Strauss et al. 2003) and during intubation (Nishino and Hiraga 1991) with laryngeal closure mediated by activity of the superior laryngeal nerve (Jafari, Prince et al. 2003). Genioglossus EMG therefore had two patterns of activity during stimulation after hemisection – first, at lower current amplitudes, an increase in EMG magnitude which remained phasic, and second, early on in stimulation at high current amplitude, very high amplitude tonic activity that was not coordinated to breathing.

The existence of multiple patterns of activity between the genioglossus and diaphragm seen here indicates that after injury, electrical stimulation at C4 was able to facilitate both ascending and descending communication between the two motor networks. However, it is unclear under which circumstances, if any, an increase in genioglossus EMG amplitude is indicative of greater respiratory drive during stimulation vs recruitment of the genioglossus for non-respiratory behaviors. Even the meaning of an increase in genioglossus EMG relative to respiratory drive is unclear, as negative airway pressure may reflexively increase genioglossus EMG in the absence of increased respiratory drive (Mathew, Abu-Osba et al. 1982, Horner, Innes et al. 1991), though increased central drive also increases EMG at a given pressure (Pillar, Fogel et al. 2001). We are unfortunately unable to determine if any of these patterns of activity are directly swallow-related since only the genioglossus was recorded in these experiments, and a further exploration of the nature of this genioglossus EMG under each circumstance should be undertaken.

### Implications for the clinic

Our lab has previously shown that inspiratory-delivered electrical stimulation using the contralesional diaphragm as a trigger can restore phasic diaphragm EMG activity (Mickle, Penaloza-Aponte et al. 2024). However, in translating this therapeutic clinically, it will be important to consider the choice of the endogenous trigger used to initiate stimulation since each endogenous signal may react differently to electrical stimulation. The potential impacts of different signal choice are highlighted here, via the differential response between the genioglossus and the contralesional diaphragm at the beginning of the open-loop stimulatory period. When using the genioglossus, a run-away positive feedback loop can occur such that stimulation drives increased genioglossus activity, disrupting the phasic respiratory patterning of genioglossus EMG output while simultaneously shutting down contralesional diaphragm EMG. As the suppression of output occurred only for the side of the diaphragm still connected to the brainstem and only when ascending spinal pathways are intact and increase genioglossus EMG output, the appropriateness of endogenous trigger may vary based on severity and level of injury.

Typically, electrical stimulation to improve swallowing and reduce dysphagia has been limited to neuromuscular electrical stimulation of swallow-related muscles. Stimulation via electrodes placed on the neck can improve swallow either by directly increasing muscular contraction via high-frequency stimulus, or by activating sensory afferents (Assoratgoon, Shiraishi et al. 2023), and has particularly been used after stroke (Poorjavad, Talebian Moghadam et al. 2014). However, these surface stimuli do not generally interact with the pattern generator for swallow. In intact rats, electrical stimulation of the thoracic spinal cord using transcutaneous electrical stimulation of the back was successful at increasing swallow-related EMG (Kitamura, Frazure et al. 2024). The increases in genioglossus EMG with stimulation after hemisection seen here hints that epidural spinal stimulation could be a viable therapeutic in improving swallow function in the injured spinal cord as well. This could be accomplished either by stimulating below the lesion if there are sufficient remaining intact ascending spinal pathways, or perhaps even stimulating above the lesion at C1 in the case of complete injury. However, extensive preclinical work focused specifically on swallow with recordings of muscles related to the pharyngeal phase during intentional swallow induction will be necessary to fully understand whether or not the increases in genioglossus EMG with stimulation are indicative of activation of swallow-related supraspinal nuclei, and if this can further translate to improvements in swallow function post-injury.

## DATA AVAILABLILITY

Data are available at https://odc-sci.org/. All analysis scripts are publicly available at https://github.com/dale-lab/ABRAP_Detangling.

## AUTHOR CONTRIBUTIONS

Alyssa Mickle: conceptualization, investigation, formal analysis software, writing – original draft; Jesús Peñaloza-Aponte: investigation, writing – reviewing & editing; Caitlin Brennan: investigation, writing – reviewing & editing; Erica A Dale: conceptualization, supervision, writing – reviewing & editing, funding acquisition, project administration

